# Establishment of CD8^+^ T cell thymic central tolerance to tissue-restricted antigen requires PD-1

**DOI:** 10.1101/2022.08.01.502412

**Authors:** Julia F. May, Rees G. Kelly, Alexander Y.W. Suen, Gina R. Rayat, Colin C. Anderson, Troy A. Baldwin

## Abstract

Highly self-reactive T cells are censored from the repertoire by both central and peripheral tolerance mechanisms upon receipt of high-affinity TCR signals. Clonal deletion is considered a major driver of central tolerance; however, other mechanisms such as induction of regulatory T cells and functional impairment have been described. An understanding of the interplay between these different central tolerance mechanisms is still lacking. We previously showed that impaired clonal deletion to a model tissue-restricted antigen (TRA) did not compromise tolerance. In this study, we determined that murine T cells that failed clonal deletion were rendered functionally impaired in the thymus. PD-1 was induced in the thymus and was required to establish cell-intrinsic tolerance to TRA in CD8^+^ thymocytes independently of clonal deletion. PD-1 signaling in developing thymocytes was sufficient for tolerance induction, however in some cases the absence of PD-L1-mediated signaling in the periphery resulted in broken tolerance at late time points. Finally, we showed that chronic exposure to high affinity antigen supported the long-term maintenance of tolerance. Taken together, our study identifies a critical role for PD-1 in establishing central tolerance in autoreactive T cells that escape clonal deletion. It also sheds light on potential mechanisms of action of anti-PD-1 pathway immune checkpoint blockade and the development of immune-related adverse events.

**Graphical Abstract:** 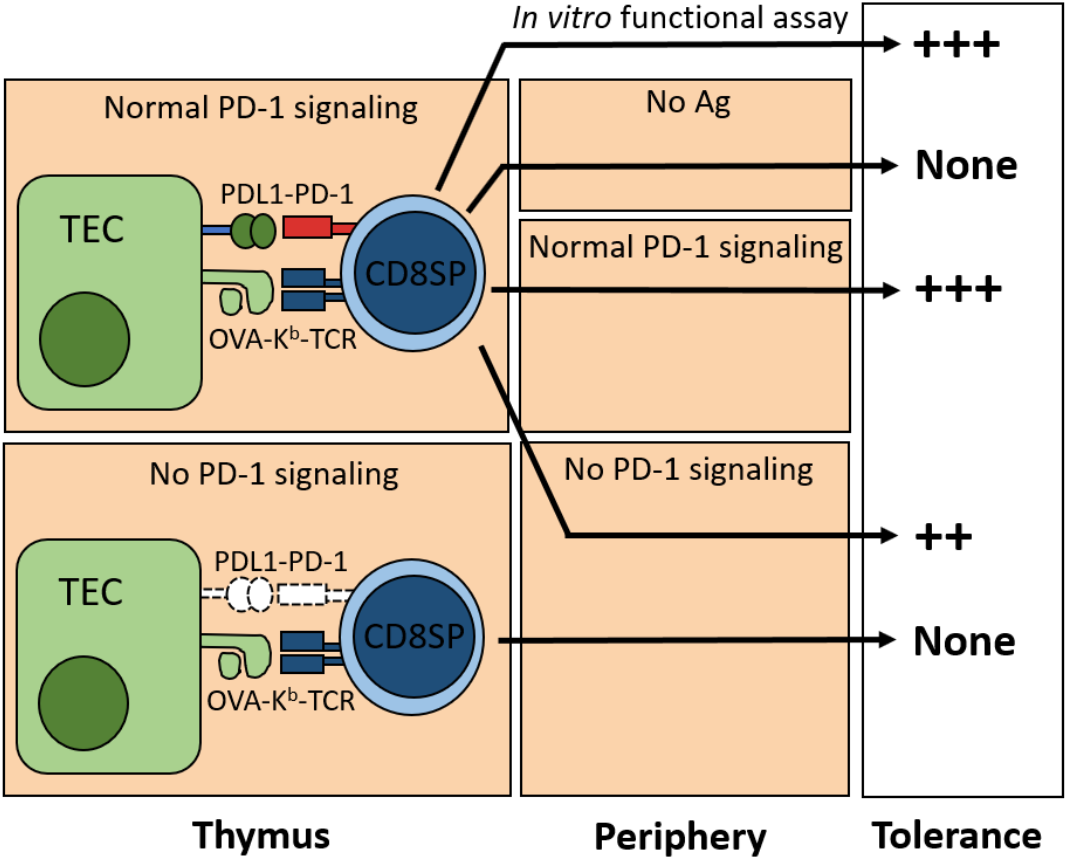

## Introduction

The outcomes of thymocyte selection shape the peripheral T cell repertoire. The affinity of the TCR for self-peptide-MHC (self-pMHC) plays a dominant role in regulating these selection outcomes, but signals derived from other receptors are also important. T cell central tolerance mechanisms include processes such as clonal deletion, anergy, and agonist selection (1). However, much is left to understand regarding the limits and interplay between these processes.

Nascent αβ TCR-committed thymocytes transition from the CD4^-^CD8^-^ double-negative (DN) stage to the CD4^+^CD8^+^ double-positive (DP) stage and acquire the ability to respond to self-pMHC by virtue of expressing a TCR. While in the thymic cortex, low affinity interactions between the αβ TCR and self-pMHC expressed by cortical thymic epithelial cells result in positive selection and further transition into CD4^-^CD8^+^ or CD4^+^CD8^-^ single-positive (SP) thymocytes (2). Conversely, strong TCR signals at the DP or SP thymocyte stage during thymic development result in several negative selection outcomes, through which autoreactive thymocytes are censored from the repertoire. Peptides from both ubiquitously expressed proteins (UbA) as well as proteins that are expressed in a tissue-restricted fashion (TRA) can induce negative selection.

Although clonal deletion, wherein autoreactive thymocytes are instructed to undergo apoptosis, has long been considered the primary central tolerance mechanism, the fate of autoreactive thymocytes that escape clonal deletion is not yet fully understood (3). Studies by Ramsdell and Fowlkes used a superantigen model of negative selection to determine that a population of superantigen-reactive CD4SP and CD8SP could escape clonal deletion but were unresponsive to superantigen stimulation (4). Similarly using TCR transgenic models, we and others recently showed that TRA-specific CD8SP and CD4SP thymocytes that encountered their high affinity antigen during development, but were unable to undergo apoptosis, continued maturation and emigrated into the periphery (5, 6) but showed functional impairment (7). More recent work using polyclonal models demonstrated that CD4SP thymocytes that encountered their high affinity antigen in the cortex were mostly deleted, but those that received a high affinity signal in the medulla were largely selected into the Treg lineage or were functionally impaired rather than being deleted (8, 9). These studies highlight the complex interplay between different central tolerance mechanisms.

Programmed cell death protein 1 (PD-1) is a co-inhibitory receptor expressed on T cells, B cells, and some subsets from the myeloid lineage (10). PD-1 ligation impairs T cell responses and thus PD-1 blockade forms the basis for some promising cancer immunotherapies by improving the function of anti-tumour CD8^+^ T cells in the tumour microenvironment, reviewed in (11). PD-1 is thought to function through its recruitment of the phosphatase SHP2, which negatively regulates the activation of TCR-proximal signaling molecules (12) and CD28-driven co-stimulation (13, 14). On lymphocytes at steady state, PD-1 expression is limited to a small fraction of thymic and peripheral cells, but is induced on thymocytes and peripheral T cells following strong TCR interactions (15, 16). The ligands for PD-1, PD-L1 and PD-L2 are expressed on the thymic epithelium and professional antigen presenting cells in the thymus and throughout the periphery (17, 18). PD-1 deficiency results in autoimmunity in pre-clinical models (19) and single nucleotide polymorphisms in PD-1 are linked to autoimmunity in humans (20–22). In the thymus, transgenic overexpression of PD-1 has been shown to modulate positive selection (23), while PD-1 deficiency in UbA-mediated models of negative selection showed little or no impact (24, 25). Studies examining a role for PD-1 in thymic tolerance induced by TRA are currently lacking.

In this study we explored the fate of MHCI-restricted thymocytes specific for a TRA in the absence of clonal deletion. We found that PD-1 was sufficient to establish cell-intrinsic tolerance in the thymus and continued PD-L1-mediated signaling in the periphery was dispensable until late time points. Additionally, chronic exposure to high affinity antigen supports the long-term maintenance of tolerance in this model. Taken together, our study identifies a role for PD-1 in establishing central tolerance in TRA-specific thymocytes that have escaped clonal deletion. Furthermore, it supports a model where cancer immunotherapies that target the PD-1 pathway act to prevent anti-tumour T cells from becoming dysfunctional in addition to rescuing them from the dysfunctional state.

## Methods

### Mice

C57BL/6j (WT, strain 000664), C57BL/6-Tg(Ins2-TFRC/OVA)296Wehi/WehiJ (RIP-mOVA, strain 005431) (26), NOD.B6-*Cd274^tm1Shr^*/J (PD-L1^-/-^, strain 018307) (27), C57BL/6-Tg(Cd8a-cre)1Itan/J (CD8-cre, strain 008766) (28), B6.Cg-*Foxn1^nu^*/J (Nude, strain 000819) (29) mice were sourced from Jackson Laboratory. C57BL/6-*Pdcd1^tm1.1Mrl^* (PD-1^fl/fl^, strain 13976) were purchased from Taconic Biosciences. The PD-L1^-/-^ mice were backcrossed to C57BL/6j mice at least 5 generations prior to intercrossing with RIP-mOVA mice. OT-I Bim^-/-^ mice were provided by Dr. Maureen McGargill (St. Jude Children’s Research Hospital, Memphis, TN, USA). PD-1^-/-^ mice were provided by Dr. Tasuku Honjo (Kyoto University, Kyoto, Japan) (30) and BTLA^-/-^ were obtained from Dr. Kenneth Murphy (Washington University, St. Louis, MO) (31). Mice with compound genotypes were intercrossed in house. Across this study, both male and female mice were used and age-matched where possible. All mice were maintained via protocols approved by the University of Alberta Animal Care and Use Committee.

### Bone marrow chimeras

T cells were depleted from donor bone marrow by I.P. administration of 100 μg anti-Thy1.2 (clone 30H12) −2 and −1 d prior to harvest. 1 d prior to transplant, recipients were irradiated with 2 doses of 450 rad, separated by a 4-hour rest period. Bone marrow was flushed from the femurs and tibias of donor mice with PBS (2 % FBS, 2 mM EDTA) and filtered through a 70 μm nylon strainer. 5×10^6^ – 10×10^6^ total BM cells were transplanted to recipients via tail vein injection. On day 1 post-transplant, 100 μg of anti-Thy1.2 antibody (clone 30H12) was administered I.P. to the recipients. For the first 4 wk post-transplant, recipient mice were given water containing Novo-Trimel (via University of Alberta Health Sciences Lab Animal Services). Chimeras were allowed to recover for at least 8 wk before further manipulation or analysis.

### Tissue collection

Mice were anaesthetized with isoflurane and euthanized via CO_2_ asphyxiation. Organs were harvested and made into single-cell suspensions by grinding through either sterile wire or nylon mesh screens in petri dishes containing RP10 media (RPMI 1640, 2.05 mM L-glutamine [Hyclone], 10 % FBS, 5 mM HEPES, 50 mM 2-mercaptoethanol, 50 μg/mL penicillin/streptomycin, 50 μg/mL gentamycin sulfate). Cells were treated with ammonium-chloride-potassium lysis buffer to lyse RBCs. Viable cell numbers were determined using Trypan Blue exclusion dye and a hemocytometer (Fisher Scientific) under light microscopy (Zeiss).

### Adoptive transfers

Whole spleen and thymus cell suspensions from adoptive transfer donors were analyzed by flow cytometry to determine the frequency of CD8^+^ Vα2^+^ (splenic) and CD8SP Vα2^+^ CD24^lo^ cells (thymic), respectively. CD8^+^ T cell enrichment was performed using the StemCell EasySep Mouse CD8^+^ T cell Isolation Kit (Cat. # 19853A) on pooled lymph nodes and spleen. The purities of isolated fractions were > 70 % in all cases. Unless otherwise stated, 5×10^6^ splenic or 4×10^6^ thymic mature CD8^+^ Vα2^+^ OT-I T cells were adoptively transferred I.V. into sub-lethally irradiated recipients (450 rad). Co-adoptive transfers involved 5×10^6^ cells per donor for a total of 10×10^6^ CD8^+^ Vα2^+^ T cells transferred.

### Cell culture and in vitro stimulation assays

Cells were cultured at 2.5×10^6^ cells/well in 48-well plates in RP10 media at 37°C in a tissue culture incubator. Whole spleen cell stimulators from C57BL/6 mice were resuspended in RP10 media at 20×10^6^ cells/mL pulsed with or without 100 nM OVA peptide sourced from AnaSpec (SIINFEKL, AS-60193-1) for 1 h at 37°C with gentle shaking every 15-20 minutes. Stimulator cells were washed three times before culture. Responder cells were either resuspended in PBS at 10×10^6^ cells/mL and stained with 1.25 μM CFSE or at 5×10^6^ cells/mL stained with 5μM CellTrace Violet. Staining was conducted for 20 min at 37°C with regular mixing before quenching with at least 4 volumes of RP10 media. Responder cells were mixed with stimulator cells at a ratio of 4:1 and incubated at 37°C for indicated time points. Proliferation and division indices were calculated using FlowJo. Proliferation index is calculated as the number of divisions within the dividing cellular population, while the division index is calculated as the average number of divisions undergone by each cell in the starting culture.

### Abs and flow cytometry

All staining was carried out in round-bottom 96 well plates using FACS buffer (PBS, 1 % FBS, 0.02 % sodium azide, 1 mM EDTA [pH 7.2]). Prior to staining, samples were pre-incubated with anti-Fc receptor blocking solution (24G.2 hybridoma supernatant) at a 1:20 dilution in FACS buffer. Surface staining was carried out at 4°C for 15 min. Antibodies were sourced from either BD Biosciences, ThermoFisher, or BioLegend. Clones used were as follows: CD4 (RM4.5), CD8 (53-6.7), CD69 (H1.2F3), PD-1 (J43), CD19 (1D3), CD25 (7D4), Vα2 (B20.1), H-2K^b^ (AF6-88.5.5.3), CD24 (M1/69), IFN-γ (XMG1.2), TNFα (MP6-XT22), CD45.1 (A20), and CD45.2 (104). Live/Dead staining was performed using LIVE/DEAD Fixable Aqua, Lime, or Blue (Thermo FisherScientific) or Fixable Viability Dye eFluor 780 (eBioscience). Biotinylated H-2K^b^-OVA monomers were obtained from the NIH tetramer facility and tetramerized in house. Intracellular staining was performed using BD Cytofix/Cytoperm Fixation/Permeabilization Solution Kit (BD Biosciences). Data was collected on either BD LSRFortessa SORP/X-20, or a Cytek Aurora spectral cytometer. Analysis was conducted using Flowjo software (TreeStar).

### Blood glucose monitoring

Blood glucose concentrations of bone marrow chimeras were monitored weekly starting at 3 wk post-transplantation using a OneTouch Ultra2 blood glucose meter and OneTouch Ultra Test Strips. Adoptive transfer recipients were monitored every 3-4 days starting 4 d post-transfer for the first month, and every week subsequently. Blood samples were acquired by tail vein bleeding. When blood glucose readings exceeded 15 mM, another reading was taken 24 h later. Mice were confirmed diabetic upon two consecutive readings above 15 mM.

### PD-1 blockade

For *in vitro* blockade experiments, cultured cells were treated continuously with 5 μg/mL of αPD-1 (clones: J43 or RMP1-14, BioXCell). For *in vivo* experiments, mice were treated I.P. with 250 μg RMP1-14 every second day for 2 wk. The blood glucose concentration of treated mice was monitored for at least 4 wk following the final antibody injection.

### Thymic transplants

Thymic lobes were isolated from 1- to 3-day old neonatal WT or RIP-mOVA mice and cultured for 7-10 days on a filter-strainer stack (filter: MF-Millipore Membrane Filter, mixed cellulose esters, hydrophilic, 0.45 μm, 13 mm, white, gridded, HAWG01300; strainer: 70 μm, nylon, FisherScientific, #22363548) in a 6 well plate containing 1.35 mM 2’-deoxyguanosine (2-dG, Sigma-Aldrich) in complete DMEM-10 (DMEM, 4.5 g/L glucose, 0.584 g/L glutamine and 3.7 g/L NaHCO_3_) to deplete cells of hematopoietic origin. Media was changed as needed. Thymi were washed for 2 h in DMEM-10 before transplantation under the kidney capsule of recipient mice following our standard protocol (32). T cell reconstitution was confirmed by the presence of CD4^+^/CD8^+^ T cells in peripheral blood 8 wk post-transplant by flow cytometry. Subsequent bone marrow transplantation was conducted 4 mo. following thymic transplant.

### Statistical analysis

Statistical analyses were conducted using Prism software (Graph-Pad). Specific tests used are listed in figure legends.

## Results

### Cell-intrinsic functional impairment is induced in antigen-specific CD8^+^ T cells in OT-I Bim^-/-^ > RIP-mOVA chimeras

We and others previously showed that Bim is required for thymic clonal deletion to tissue-restricted antigen (TRA) using the OT-I TCR transgenic system (5–7). OT-I Bim^-/-^ T cells that escaped thymic clonal deletion in RIP-mOVA recipients did not go on to cause autoimmunity due to functional impairments characterized by defects in activation, proliferation, and cytokine production. The mechanism(s) behind this impairment was unclear but may be due to cell-extrinsic suppression or cell-intrinsic anergy (33–35). We hypothesized that if a population of dominant suppressors existed in OT-I Bim^-/-^ > RIP-mOVA chimeras, they should suppress the activation and proliferation of wildtype OT-I T cells. CFSE labelled splenocytes from CD45.1^+^ OT-I mice were mixed at a 1:1 ratio with splenocytes from OT-I Bim^-/-^ > RIP-mOVA chimeras and stimulated with OVA peptide-pulsed splenocytes *in vitro*. General gating strategies are outlined in Fig. S1. Activation marker induction (Fig. 1A, B) and proliferation measured by proliferation index (Fig. 1C-D, Fig. S2A) in WT OT-I T cells was not significantly reduced by co-culture with splenocytes from OT-I Bim^-/-^ > RIP-mOVA chimeras. As we previously found, on day 2 post-stimulation, the induction of activation markers CD69 and CD25 on OT-I splenocytes from OT-I Bim^-/-^ > RIP-mOVA chimeras was diminished compared to that of splenocytes from wildtype (WT) OT-I CD45.1^+^ mice.

**Figure 1:**
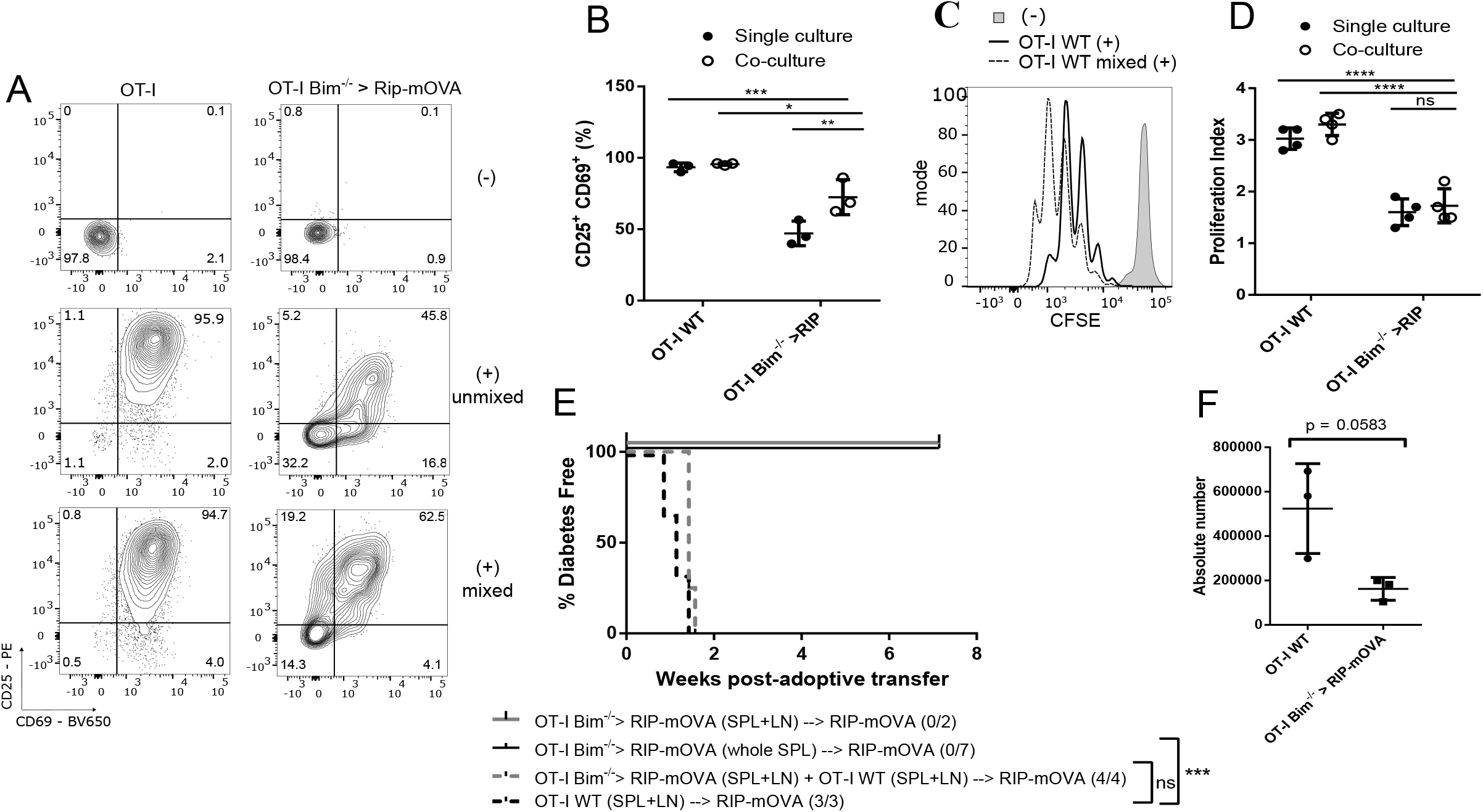
TRA-Specific T Cells in OT-I Bim^-/-^ > RIP-mOVA Chimeras Are Functionally Impaired Via Cell-Intrinsic Mechanisms. A, Splenocytes from OT-I WT (CD45.1), OT-I Bim^-/-^ > RIP-mOVA or a 1:1 mixture of the above splenocytes were cultured with un-pulsed stimulator splenocytes or stimulator splenocytes pulsed with OVA peptide in vitro. Activation was measured by CD69 and CD25 upregulation on day 2. B, Compilation of data from A (n = 3). C, Day 3 proliferation of cultures described in A measured by dilution of CFSE dye. D, Compilation of proliferation indices from cultures described in A (n = 4). E, Incidence of diabetes in sub-lethally irradiated RIP-mOVA recipients of adoptively transferred splenic CD8^+^ Vα2^+^ OT-I T cells from OT-I Bim^-/-^ > RIP-mOVA and/or OT-I WT. Note that 7 recipients with OT-I Bim^-/-^ > RIP-mOVA transfers received whole spleen while all other recipients in this experiment received CD8-enriched isolates from pooled spleen and lymph nodes (SPL+LN). Log-rank (Mantel-Cox) test results are relative to the OT-I WT → RIP-mOVA group. Data are representative of 4 separate cohorts. F, Recipients of CD8-enriched isolates (SPL+LN) from OT-I Bim^-/-^ > RIP-mOVA and OT-I WT described in E were analyzed for expansion of either donor cell population within the spleen. Total donor cells were identified by OVA-K^b^ tetramer staining. For each assay described above, the number of independent experiments (IE) ≥ 3. Data are mean ± SD. Asterisks represent statistical significance as determined by a two-way ANOVA with Sidak’s multiple comparisons test (B, D), a paired t test (F) or Log-rank test (E). Statistical comparisons between other groups of interest are indicated. * p ≤ 0.05, ** p ≤ 0.01, *** p ≤ 0.001, **** p ≤ 0.0001.

To assay functional impairment *in vivo*, we co-adoptively transferred either whole spleen or enriched CD8^+^ T cells from OT-I Bim^-/-^ > RIP-mOVA chimeras with splenocytes from CD45.1^+^ OT-I WT mice (1:1 ratio) into sub-lethally irradiated RIP-mOVA recipients. Neither whole spleen nor enriched CD8^+^ OT-I T cells from OT-I Bim^-/-^ > RIP-mOVA chimeras alone were diabetogenic, suggesting that a dominant suppressor population of non-T cell origin was not responsible for functional impairment in these chimeras (Fig. 1E). Enriched CD8^+^ OT-I T cells from OT-I Bim^-/-^ > RIP-mOVA chimeras co-adoptively transferred with WT OT-I T cells did not have a dominant suppressive effect on diabetogenicity (Fig. 1E). In these co-adoptive transfer recipients, WT OT-I T cells also expanded to a greater extent than those from OT-I Bim^-/-^ RIP-mOVA chimeras (Fig. 1F). Functional impairment of OT-I Bim^-/-^ T cells from RIP-mOVA chimeras was also demonstrated by a reduced capacity for lymphopenia-induced proliferation compared to control cells from OT-I Bim^-/-^ > WT chimeras (Fig. S2B). Collectively, these data suggest that a population of Ag-specific suppressors are not generated in OT-I Bim^-/-^ > RIP-mOVA mice and that functional impairment of OT-I Bim^-/-^ T cells from these chimeras is cell-intrinsic.

### TRA-specific T Cells in OT-I Bim^-/-^ >RIP-mOVA chimeras are rendered functionally impaired in the thymus

After determining that the functional impairment of OT-I T cells from OT-I Bim^-/-^ > RIP-mOVA chimeras was cell-intrinsic, we investigated whether this state was established during development in the thymus or in the periphery. The thymus is the initial site of high affinity antigen encounter and induction of non-responsiveness in thymocytes has been reported previously (36). We stimulated CFSE-labelled thymocytes from OT-I Bim^-/-^ > WT and OT-I Bim^-/-^ > RIP-mOVA chimeras with OVA peptide pulsed splenocytes and examined activation and proliferation. On day 2, expression of activation markers CD69 and CD25 was impaired in OT-I Bim^-/-^ > RIP-mOVA thymocytes compared to OT-I Bim^-/-^ > WT controls (Fig. 2B). Additionally on day 3, thymocytes from OT-I Bim^-/-^ > RIP-mOVA chimeras exhibited a defect in proliferation compared to OT-I Bim^-/-^ > WT thymocytes (Fig. 2C-D). These impaired responses were not likely to be due to a reduction in more mature CD8SP thymocyte subsets in OT-I Bim^-/-^ > RIP-mOVA recipients since the thymi of these mice demonstrated enhanced frequencies of mature (M2; CD69^lo^ H-2K^b hi^) thymocytes over WT controls (Fig. 2E).

**Figure 2:**
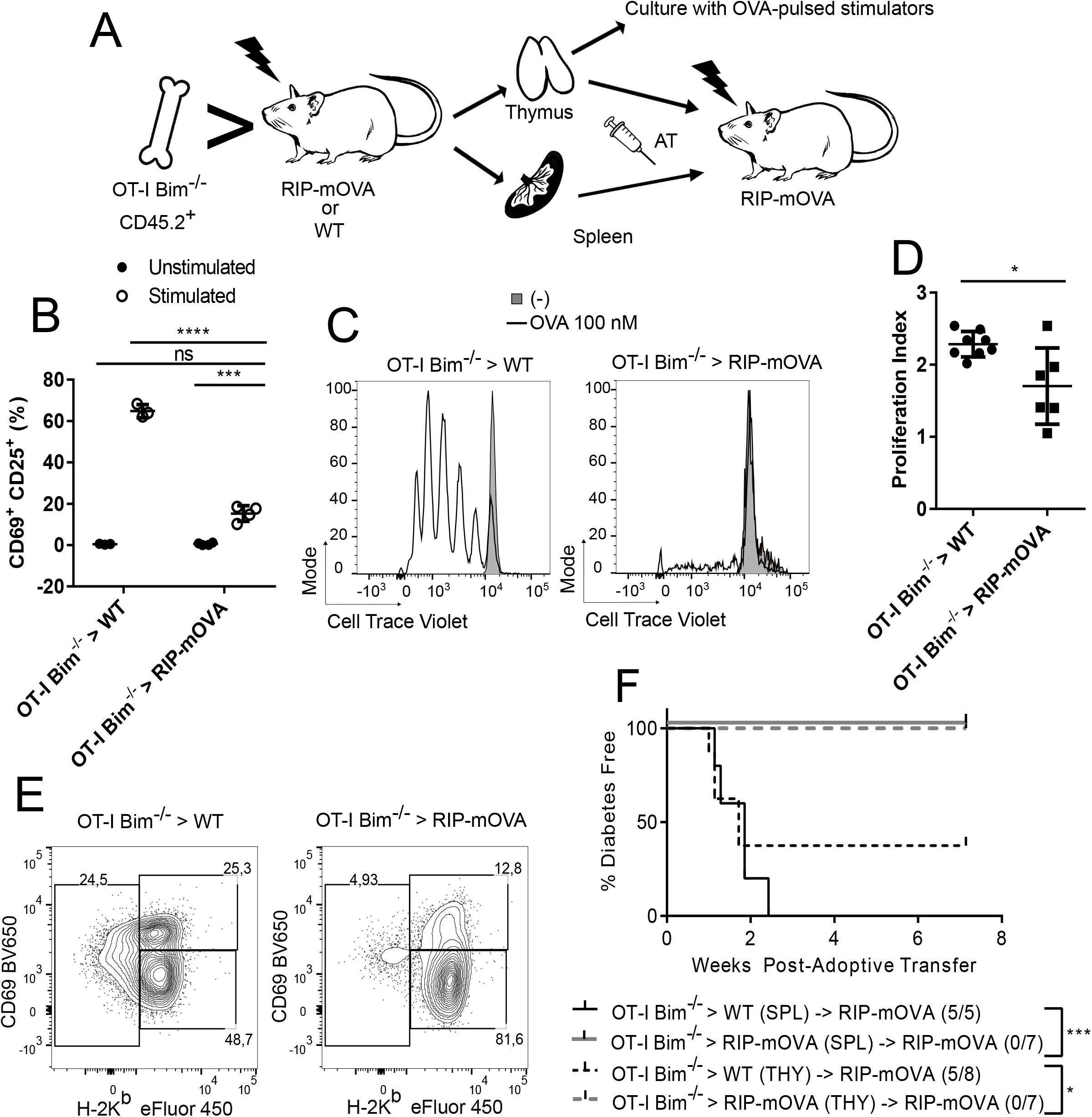
Splenic and Thymic TRA-Specific T Cells in OT-I Bim^-/-^ > RIP-mOVA Chimeras Are Functionally Impaired. A, Schematic representing experiments carried out in B-F. Activation marker induction (B), representative histograms for thymocyte proliferation (C), and compiled proliferation indices (D) of OT-I Bim^-/-^ > RIP-mOVA (n = 4) or OT-I Bim^-/-^ > WT thymocytes (n = 3) following *in vitro* stimulation. Open histograms represent culture with OVA peptide-pulsed stimulators, while greyed histograms represent no-peptide control stimulators. E, Distribution of Vα2^+^ CD8SP OT-I thymocytes among semi-mature (SM; H-2K^b lo^), Mature 1 (M1; CD69^hi^ H-2K^b hi^), and Mature 2 (M2; CD69^lo^ H-2K^b hi^) population. F, From either OT-I Bim^-/-^ > RIP-mOVA or > WT bone marrow chimeras, 2-5×10^6^ Vα2^+^ CD24^lo^ thymocytes (THY) or Vα2^+^ OT-I T splenocytes (SPL) from whole spleen were transferred to sub-lethally irradiated RIP-mOVA recipients, which were then monitored for the induction of diabetes. Log-rank (Mantel-Cox) test results are relative to organ transfer groups. For each assay described above, the number of independent experiments (IE) ≥ 3. Data are mean ± SD. Asterisks represent statistical significance as determined by a two-way ANOVA with Sidak’s multiple comparisons test (B), unpaired t test with Welch’s correction (D), or Log-rank test (E). Statistical comparisons between other groups of interest are indicated. * p ≤ 0.05, ** p ≤0.01, *** p ≤ 0.001, **** p ≤ 0.0001.

To determine if the observed *in vitro* defects in activation and proliferation translated into functional *in vivo* impairment, we adoptively transferred either thymocytes or splenocytes from OT-I Bim^-/-^ > RIP-mOVA or OT-I Bim^-/-^ > WT chimeras into sub-lethally irradiated RIP-mOVA recipients to assess diabetogenicity. Most recipients of thymocytes and all recipients of splenocytes from OT-I Bim^-/-^ > WT chimeras developed diabetes as expected. In contrast, none of the sub-lethally irradiated RIP-mOVA recipients that received thymocytes or splenocytes from OT-I Bim^-/-^ > RIP-mOVA chimeras became diabetic (Fig. 2F). Collectively, these data suggest that functional impairment of OT-I Bim^-/-^ > RIP-mOVA T cells is established in the thymus.

### PD-1 signals are required for T cell tolerance in OT-I > RIP-mOVA chimeras

The functional impairment seen in OT-I Bim^-/-^ > RIP-mOVA thymocytes and splenocytes may be driven by co-inhibitory molecules and/or other factors that inhibit T cell activity. As such, we investigated whether various known co-inhibitors were differentially expressed in OT-I > RIP-mOVA thymocytes and splenocytes that escaped clonal deletion compared to OT-I > WT controls. We found enhanced PD-1 expression on OT-I T cells from both the thymus and periphery of OT-I > RIP-mOVA (Fig. 3A) and OT-I Bim^-/-^ > RIP-mOVA chimeras compared to OT-I cells from WT recipients, while the expression of other co-inhibitory molecules was not appreciably altered (Fig. S3A). Given the increased PD-1 expression, and that previous literature supports a role for PD-1 in enforcing tolerance (10), we hypothesized that PD-1 was playing an active role in establishing and/or maintaining the functional impairment seen in TRA-specific T cells that escaped clonal deletion in OT-I Bim^-/-^ > RIP-mOVA chimeras.

**Figure 3:**
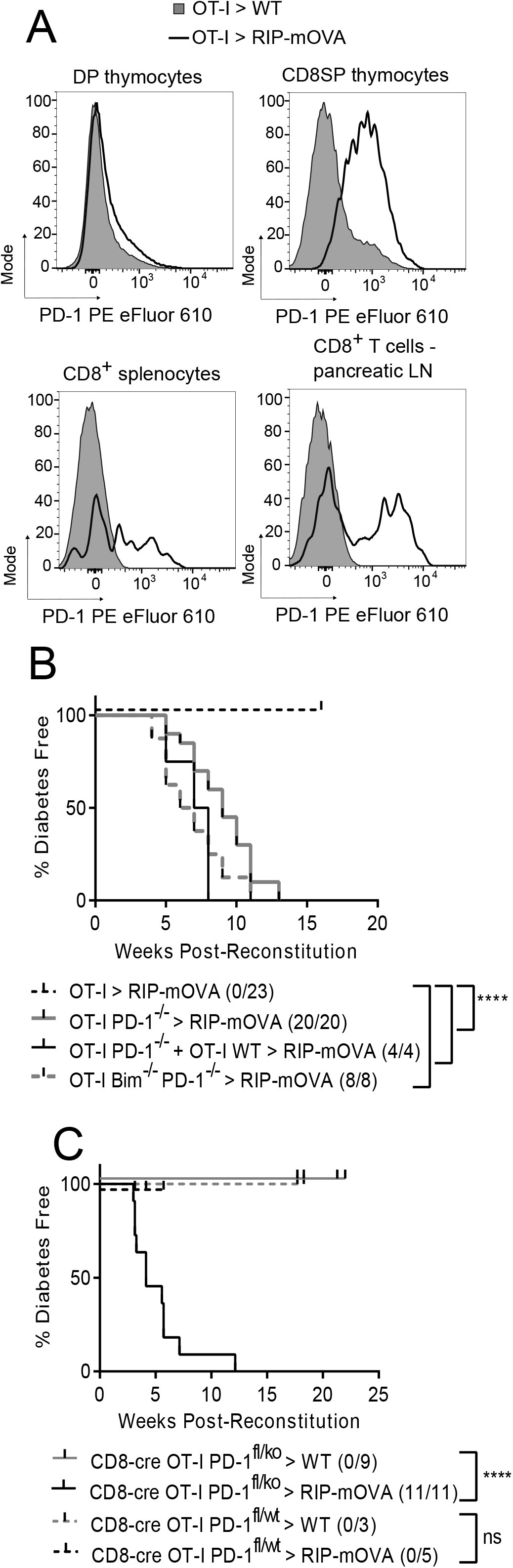
PD-1 signals in the T cell compartment necessarily support tolerance to TRA. A, PD-1 expression on OT-I T cells across organs in OT-I > RIP-mOVA (open histogram) and OT-I > WT (greyed histogram) bone marrow chimeras. Data is representative of n ≥ 3. B, Incidence of diabetes in RIP-mOVA chimeras with OT-I WT, OT-I PD-1^-/-^, a 1:1 mix of the former and latter, and OT-I Bim^-/-^ PD-1^-/-^ bone marrow. Data are generated from 3 separate cohorts of chimeras. C, Incidence of diabetes in RIP-mOVA or WT chimeras with CD8-cre OT-I PD-1^fl/ko^ or PD-1^fl/WT^ bone marrow; PD-1^fl/WT^ > RIP-mOVA chimeras were sacrificed alongside their PD-1^fl/ko^ counterparts to confirm the timing of PD-1 ablation (IE = 4). Data are compiled from 4 separate chimera cohorts. Asterisks represent statistical significance as determined by Log-rank test. * p ≤ 0.05, ** p ≤ 0.01, *** p ≤ 0.001, **** p ≤ 0.0001.

We intercrossed OT-I and PD-1^-/-^ mice to generate OT-I PD-1^-/-^ mice to determine if PD-1 regulated tolerance in OT-I > RIP-mOVA chimeras. Bone marrow (BM) chimeras using OT-I WT or OT-I PD-1^-/-^ BM and either WT or RIP-mOVA recipients were generated. We observed a 100% incidence of diabetes in OT-I PD-1^-/-^ > RIP-mOVA chimeras, compared to 0% incidence in OT-I WT > RIP-mOVA controls (Fig. 3B). Next, we investigated if autoimmunity in OT-I PD-1^-/-^ > RIP-mOVA chimeras was dependent on all donor OT-I T cells being PD-1^-/-^. We generated mixed bone marrow chimeras with a 1:1 ratio of OT-I WT to OT-I PD-1^-/-^ BM and similarly found that 100% of mixed OT-I WT/OT-I PD-1^-/-^ > RIP-mOVA chimeras developed diabetes (Fig. 3B). These data suggested that only a fraction of the overall pool of OT-I T cells needed to be PD-1^-/-^ to induce autoimmunity. This finding further supports the case that functional impairment of OT-I T cells in OT-I Bim^-/-^ > RIP-mOVA was cell intrinsic and not due to a population of dominant suppressors. To ensure that the absence of Bim itself was not preventing autoimmunity, we generated OT-I PD-1^-/-^ Bim^-/-^ > RIP-mOVA chimeras and again observed a 100% incidence of diabetes, suggesting that PD-1 signals with or without the presence of Bim are necessary for tolerance.

To further solidify that the deletion of PD-1 specifically and not any co-inhibitory receptor impairs tolerance in this context, we investigated whether deletion of B and T lymphocyte attenuator (BTLA) impaired tolerance. BTLA is a co-inhibitory receptor expressed on thymocytes following positive selection (37), and has been thought to play a similar role to PD-1 in regulating T cell tolerance (31). We generated OT-I BTLA^-/-^ > WT or RIP-mOVA chimeras and found that none of the OT-I BTLA^-/-^ > RIP-mOVA chimeras developed diabetes, suggesting that elimination of any co-inhibitory receptor does not impair tolerance (Fig. S4).

Finally, to determine whether PD-1 signaling specifically within the T cell compartment was necessary to induce tolerance, we conditionally deleted PD-1 in CD8^+^ T cells using *pdcd1* floxed mice. CD8-Cre mice were intercrossed with OT-I, constitutive PD-1^-/-^, and *pdcd1* floxed mice to generate OT-I CD8-Cre PD-1^fl/ko^ mice. In OT-I CD8-Cre PD-1^fl/ko^ > RIP-mOVA chimeras, the loss of PD-1 became evident at the SM-M1 CD8SP thymocyte stages (Fig. S3B) and all OT-I CD8-Cre PD-1^fl/ko^ > RIP-mOVA chimeras experienced rapid onset of diabetes while OT-I CD8-Cre PD-1^fl/wt^ > RIP-mOVA chimeras were protected (Fig. 3C). Therefore, PD-1 signaling specifically within the CD8^+^ T cell compartment is required to establish tolerance in this model. Here and in subsequent diabetes analyses, we euthanized mice from some groups early for the purpose of *ex vivo* analysis (indicated by tick marks). Thus, mice from some groups were not followed to the end of the study, and therefore in some cases our reported incidence of diabetes may be underestimates. The Kaplan-Meier graphs depicting the percentage of mice remaining diabetes-free is accurate as the data points censored early are not included in this calculation.

### OT-I PD-1^-/-^ > RIP-mOVA chimeras exhibit a mild impairment in clonal deletion and increased numbers of OT-I CD8^+^ T Cells in the periphery

Because 100% of the OT-I PD-1^-/-^ > RIP-mOVA chimeras developed autoimmune diabetes, we next investigated the mechanism by which PD-1 signals contribute to tolerance to TRA. We first determined if PD-1 modulated clonal deletion of OT-I thymocytes after high-affinity antigen encounter in the thymus of RIP-mOVA recipients. We examined the CD4 by CD8 profile of thymocytes from OT-I WT > RIP-mOVA and OT-I PD-1^-/-^ > RIP-mOVA chimeras and observed a modest reduction in the frequency of CD8SP thymocytes and a similar CD8 dulling in this compartment compared to OT-I WT > RIP-mOVA controls (Fig. 4A, Left). Furthermore, we observed a significant reduction in the frequency and total number of mature (Va2^+^ CD24^lo^) CD8SP thymocytes in both OT-I WT and OT-I PD-1^-/-^ > RIP-mOVA chimeras compared to WT recipient controls, indicating intact clonal deletion in the absence of PD-1 (Fig. 4A, Right; B). We did observe a small increase in the number of mature CD8SP thymocytes from OT-I PD-1^-/-^ > RIP-mOVA chimeras compared to OT-I > RIP-mOVA controls, but this was not statistically significant (Fig. 4B). Overall, these data suggest that while the vast majority of OT-I thymocytes encountered cognate antigen and underwent clonal deletion in OT-I PD-1^-/-^ > RIP-mOVA recipients, the remaining PD-1-deficient OT-I T cells in these chimeras still went on to induce autoimmune diabetes.

**Figure 4:**
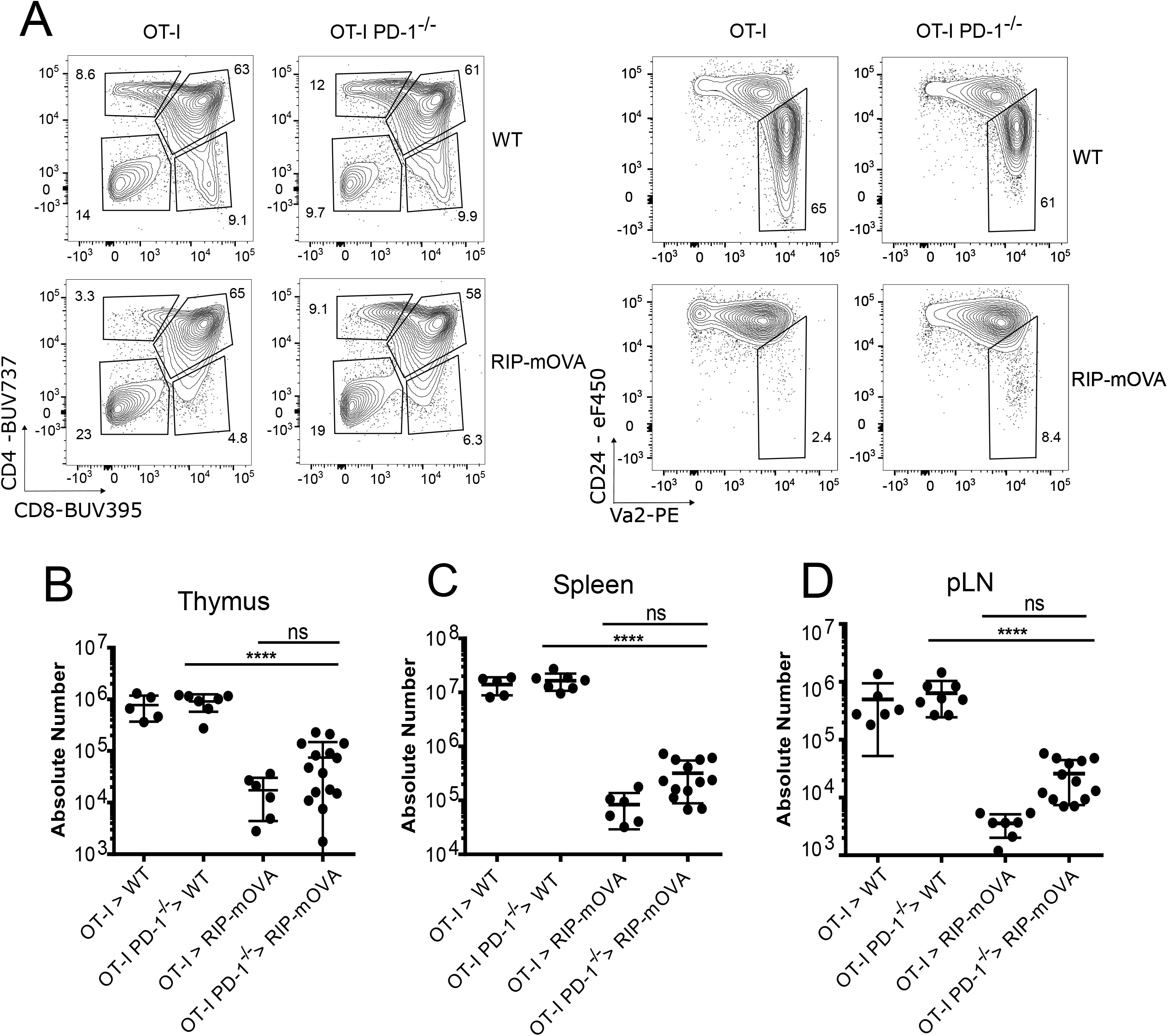
OT-I PD-1^-/-^ > RIP-mOVA chimeras exhibit a mild impairment in clonal deletion and increased numbers of OT-I Vα2^+^ CD8^+^ T cells in the periphery. A, (Left) Thymic profiles of WT or PD-1^-/-^ OT-I > RIP or B6 chimeras; (Right) Frequency of mature thymocytes within the CD8SP subset. B, Absolute number of CD8^+^ Vα2^+^ CD24^10^ mature OT-I thymocytes in chimeras from A. A data point from the PD-1^-/-^ > B6 group was determined to be an outlier using the Grubb’s test (p < 0.05) and was therefore omitted. C, Absolute number of CD8^+^ splenic OT-I. D, Absolute number of CD8^+^ OT-I in the pancreatic lymph nodes. Data above represent ≥ 5 independent experiments (IE). Data are mean ± SD. Asterisks represent statistical significance as determined by a two-way ANOVA with Sidak’s multiple comparisons test. * p ≤ 0.05, ** p ≤ 0.01, *** p ≤ 0.001, **** p ≤ 0.0001.

We next sought to determine if PD-1 deficiency had an impact on the number of peripheral OT-I CD8^+^ T cells in OT-I PD-1^-/-^ > RIP-mOVA chimeras compared to OT-I WT > RIP-mOVA controls. With both OT-I WT and OT-I PD-1^-/-^ BM, the number of Va2^+^ CD8^+^ T cells in the spleen and pancreatic lymph nodes demonstrated a significant overall reduction in RIP-mOVA chimeras compared to WT recipient controls, again suggesting intact clonal deletion. Like what was observed in the thymus, there was an increase in the number of OT-I T cells in the spleen and pancreatic LNs from OT-I PD-1^-/-^ > RIP-mOVA chimeras compared to OT-I WT > RIP-mOVA controls, but this was not statistically significant (Fig. 4C, D). Overall, in this model, clonal deletion appears largely intact in the absence of PD-1. Thus, we conclude PD-1 contributes to tolerance in this context by a mechanism distinct from thymic clonal deletion.

### Interruption of PD-1:PD-L1 interactions does not reverse established tolerance in peripheral or thymic OT-I T cells

We next investigated the specific requirements for the PD-1-driven tolerance induced in OT-I Bim^-/-^ > RIP-mOVA chimeras. Given that OT-I T cells from OT-I Bim^-/-^ > RIP-mOVA chimeras expressed PD-1 and are ostensibly impaired in a cell-intrinsic manner, we asked whether PD-1-dependent tolerance required continued PD-1 signaling or was stable even when PD-1 signaling was withdrawn. To address this question, we stimulated splenocytes from OT-I Bim^-/-^ > RIP-mOVA chimeras with OVA-peptide pulsed splenocytes *in vitro* with or without well-characterized blocking anti-PD-1 monoclonal antibodies. On day 2, we observed that PD-1 blockade did not result in an increase in CD69 and CD25 expression on OT-I Bim^-/-^ splenocytes (Fig. 5A) or thymocytes (Fig. 5C) from RIP-mOVA recipients. Additionally, the proliferation index of Va2^+^ CD8^+^ splenocytes from OT-I Bim^-/-^ > RIP-mOVA chimeras was not altered by PD-1 blockade on day 3 (Fig. 5B). Similarly, *in vivo* administration of anti-PD-1 blocking antibodies to OT-I > RIP-mOVA or OT-I Bim^-/-^ > RIP-mOVA bone marrow chimeras failed to break tolerance, suggesting tolerance was stable even when PD-1 signaling was presumably interrupted (Fig. 5D).

**Figure 5:**
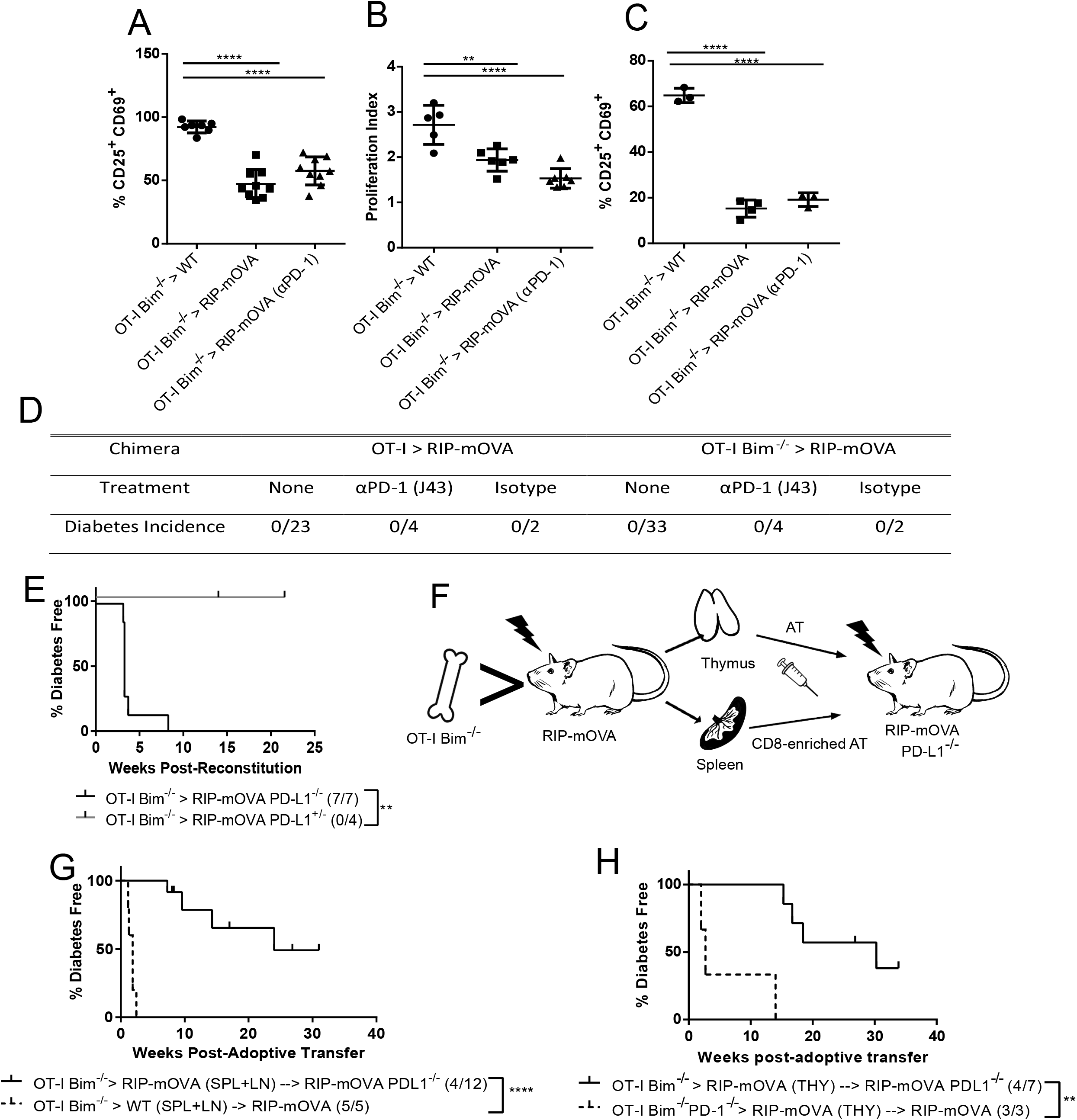
Acute PD-1 signaling within the thymic compartment is sufficient to induce tolerance to TRA. Splenocytes from either OT-I Bim^-/-^ > RIP or WT chimeras were cultured *in vitro* with peptide pulsed whole-spleen stimulators +/- αPD-1 antagonist antibody. A, Day 2 activation marker expression for responding OT-I splenocytes (stimulated groups shown). B, Compilation of day 3 proliferation indices for responding OT-I splenocytes. A data point in RIP-mOVA (aPD-1, proliferation index) group was determined to be an outlier by the Grubb’s test (p < 0.05) and removed from the dataset. C, Day 2 activation marker expression for OT-I thymocytes stimulated as in A (n ≥ 3). D, Incidence of diabetes in chimeras treated with αPD-1 antagonist antibody or isotype control. E, Incidence of diabetes in OT-I Bim^-/-^ > RIP-mOVA PD-L1^-/-^ chimeras. The number of independent experiments ≥ 3. F, Schematic demonstrating experiments conducted in G-H. G, Incidence of diabetes in sub-lethally irradiated RIP-mOVA PD-L1^-/-^ recipients of adoptively transferred CD8-enriched CD8^+^ Vα2^+^ splenocytes from OT-I Bim^-/-^ > RIP-mOVA chimeras. H, Incidence of diabetes in sub-lethally irradiated RIP-mOVA PD-L1^-/-^recipients of adoptively transferred thymocytes from OT-I Bim^-/-^ > RIP-mOVA chimeras. For each assay described above, the number of independent experiments (IE) ≥ 4 unless otherwise stated. Data are mean ± SD. Asterisks represent statistical significance as determined by one-way ANOVA with Sidak’s multiple comparisons tests (A-C) or Log-rank tests (E, G, H). * p ≤ 0.05, ** p ≤ 0.01, *** p ≤ 0.001, **** p ≤ 0.0001.

Given some potential limitations with the use of antibody-mediated blockade, we moved to an adoptive transfer approach with PD-L1^-/-^ recipients. Since PD-L2 is another ligand for PD-1, we first determined if PD-L2 could compensate for the loss of PD-L1 and enforce tolerance in PD-L1^-/-^ recipients. We generated OT-I Bim^-/-^ > RIP-mOVA PD-L1^-/-^ bone marrow chimeras and found these chimeras rapidly became diabetic, while those heterozygous for PD-L1 did not, suggesting PD-L2 could not compensate for the loss of PD-L1 in inducing or maintaining tolerance in this setting (Fig. 5E). We then adoptively transferred splenocytes from OT-I Bim^-/-^ > RIP-mOVA chimeras into sub-lethally irradiated RIP-mOVA PD-L1^-/-^ recipients to cease PD-L1-dependent signaling (detailed in Fig. 5F). In contrast to non-tolerant controls, most RIP-mOVA PD-L1^-/-^ recipients that received OT-I Bim^-/-^ > RIP-mOVA splenocytes did not become diabetic, and in the recipients in which diabetes was observed, disease was late-onset (Fig. 5G). 4 of the 12 recipients for this group were removed from the study at 8 weeks (indicated by tick-marks); while they may have become diabetic at later time points, they nevertheless remained healthy past the time where all positive control mice became diabetic. This suggests that a continuous PD-1 signal mediated by PD-L1 is not required for the maintenance of tolerance, but in the absence of PD-L1-mediated signals, tolerance wanes over extended periods of time.

Since functional impairment in tolerant chimeras is observable in thymocytes, we hypothesized that PD-1 signaling within the thymic compartment may be sufficient to establish tolerance in this model. To address this, we collected thymocytes from OT-I Bim^-/-^ > RIP-mOVA chimeras and transferred them into sub-lethally irradiated RIP-mOVA PD-L1^-/-^ recipients so that PD-1 signaling would be limited to the thymus. Like the splenocyte adoptive transfer, transferred thymocytes remained tolerant for an extended period in contrast to non-tolerant controls, with tolerance waning at late time points in some cases (Fig. 5H). These data indicate that acute PD-1 signaling in the thymus is critically important to the establishment of tolerance, but continued PD-L1-dependent signals only support the maintenance of tolerance at late time points.

### Continuous exposure to antigen in the periphery is required to maintain functional impairments in OT-I Bim^-/-^ >RIP-mOVA chimeras

With our finding that acute PD-1 signaling in the thymus was sufficient to establish tolerance, but dispensable for the initial maintenance of tolerance in the periphery, we wondered whether continued exposure to high affinity antigen in the periphery was required to maintain tolerance. To tackle this question, we transplanted either a WT or RIP-mOVA neonatal thymus under the kidney capsule of nude (athymic) recipients, and at least 8 weeks later lethally-irradiated the thymic transplant recipients and transplanted OT-I Bim^-/-^ donor bone marrow. In recipients of a RIP-mOVA thymus, exposure to high affinity OVA is limited to the thymus and withdrawn from OT-I CD8^+^ cells upon emigration from the thymus (Fig. 6A).

**Figure 6:**
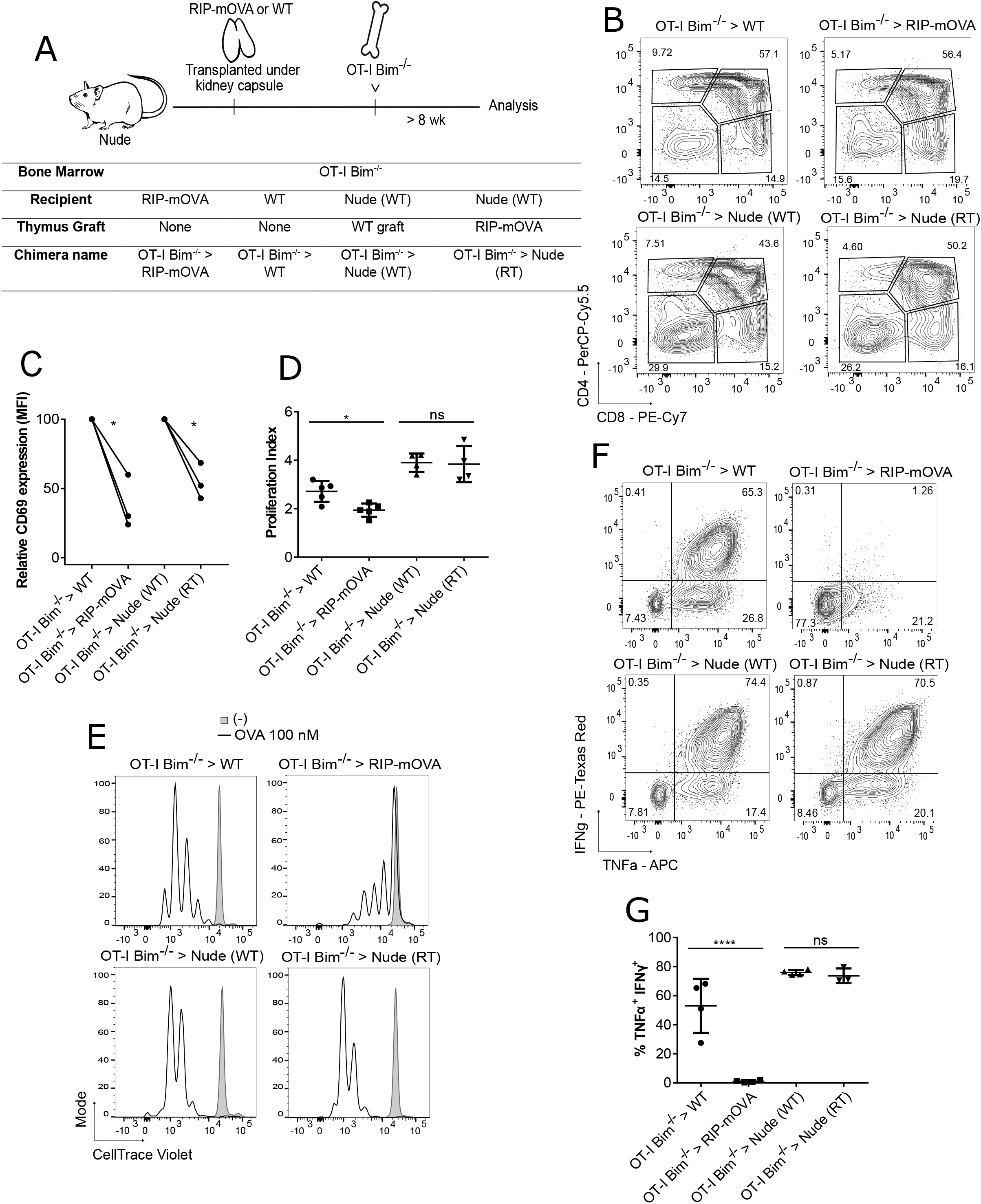
Continuous Exposure to Antigen in the Periphery is Required to Maintain Functional Impairments in OT-I Bim^-/-^ > RIP-mOVA Chimeras. A, Schematic showing experiments conducted in B-G. OT-I Bim^-/-^ bone marrow was transferred into lethally-irradiated WT or RIP-mOVA recipients, or nude recipients. Nude recipients were previously grafted with either a WT or RIP-mOVA thymus under the kidney capsule. B, Thymic CD8 by CD4 profile for each chimera described in A. C, Thymocytes from the chimeras described in A were cultured *in vitro* with peptide-pulsed whole-spleen cell stimulators and analyzed on day 2 for activation marker expression (n = 3); values are adjusted relative to WT or Nude (WT) controls within independent experiments. D, Splenocytes from the chimeras described in A were stimulated as in C. *In vitro* cultures were analyzed on day 3 for proliferation. E, Representative histograms of proliferation data described in D. Open histogram represents stimulated treatment (peptide pulsed-stimulators), greyed histogram represents no-peptide control stimulators; OT-I Bim^-/-^ > WT (n = 5), OT-I Bim^-/-^ > RIP-mOVA (n = 5), OT-I Bim^-/-^ > Nude (WT) (n = 4), OT-I Bim^-/-^ > Nude (RT) (n = 4). F, *In vitro* cultures were re-stimulated for 4 hours on day 5 in the presence of Brefeldin A and analyzed for TNFα and IFNγ production. G, Compilation of data described in F; OT-I Bim^-/-^ > WT (n = 4), OT-I Bim^-/-^ > RIP-mOVA (n = 4), OT-I Bim^-/-^ > Nude (WT) (n = 4). For the OT-I Bim^-/-^ > Nude (RT) group, only 3 replicates are listed; a fourth data point was determined to be an outlier with Grubb’s test (p < 0.05) and was thus omitted. For each assay described above, the number of independent experiments (IE) ≥ 3. Data are mean ± SD. Asterisks represent statistical significance as determined by a paired t test (C) or one-way ANOVA with Sidak’s multiple comparisons tests (D, G). * p ≤ 0.05, ** p ≤ 0.01, *** p ≤ 0.001, **** p ≤ 0.0001.

We harvested the transplanted thymus from both groups and examined the CD4/CD8 profile of OT-I thymocytes. We observed similar frequencies of CD8SP thymocytes and CD8 dulling between the thymus transplant recipients and the corresponding chimeric controls (Fig. 6B). Following *in vitro* stimulation, thymocytes from these groups also demonstrated equivalent functional impairments as assessed by CD69 induction on day 2 post-activation (Fig. 6C). Additionally, splenocytes from OT-I Bim^-/-^ > RIP-mOVA chimeras showed diminished proliferation and cytokine production (TNFα and IFNγ) compared to the corresponding WT control following *in vitro* stimulation (Fig. 65D-G). However, in OT-I Bim^-/-^ > Nude (RT) chimeras, proliferation and cytokine production were not significantly different from the OT-I Bim^-/-^ > Nude (WT) control, suggesting restored function in OT-I T cells when exposure to high affinity antigen is withdrawn after thymic egress (Fig. 6D-G). Overall, it appears that, in contrast to PD-1 signaling, persistent exposure to high affinity antigen is required to maintain the functional impairments we see in OT-I Bim^-/-^ > RIP-mOVA chimeras.

## Discussion

Clonal deletion of strongly self-reactive thymocytes is a well-described mechanism of T cell central tolerance, however, potent non-deletional mechanisms also exist. Previously, we and others showed that while Bim is required for clonal deletion to TRA in OT-I Bim^-/-^ > RIP-mOVA bone marrow chimeras (5), it is dispensable for tolerance (6, 7). In the present study, we demonstrated that TRA-specific T cells that escape clonal deletion in these chimeras are rendered functionally impaired in a cell-intrinsic manner during antigen encounter in the thymus. PD-1 expression during negative selection in the thymus was required to establish tolerance, but continued PD-1 signaling in the periphery only contributed to the maintenance of this tolerance at later time points. Additionally, we found antigen persistence in the periphery was needed to maintain a state of tolerance, as has previously been reported (38–41). Collectively, these data highlight the critical window during which PD-1 signaling results in the establishment of tolerance and shed light on potential mechanisms by which PD-1 blockade functions in the context of cancer immunotherapy and development of immune-related adverse events.

While the removal of autoreactive T cell clones via thymic clonal deletion has long been considered a primary mechanism for establishing T cell tolerance, conventional T cells that recognize self-peptides with high affinity can be readily found in healthy individuals (42). Like in the case of regulatory T cells, some of these T cells are actively selected for, while in other cases these T cells may be products of failed clonal deletion. By deleting Bim and blocking clonal deletion, a large population of highly self-reactive T cells are released into the periphery, yet tolerance is maintained (6, 7). Similarly, recent studies in polyclonal mice have revealed limited deletion of TRA-specific T cells; rather, tolerance was enforced by Treg-mediated suppression (8), or T cell-intrinsic functional impairment (9, 43). Indeed, we and others previously observed an increased number and frequency of anergic-phenotype CD4^+^ thymocytes and splenocytes when Bim was absent (44, 45). We find that functional impairment is observed both in thymocytes and peripheral T cells suggesting tolerance is first established in the thymus. These data are consistent with the recent findings of Malhotra *et al*. where MHCII-restricted thymocytes specific for a TRA showed reduced responsiveness (9). Similar findings were reported for thymocytes in an allo-reactive MHCI-restricted TCR transgenic model (36) and in a super-antigen mediated model of tolerance (4).

In previous studies where thymocytes were found to be functionally impaired, it was not determined if the impairment was due to cell-extrinsic dominant suppression or through cell-intrinsic mechanisms. In addition to CD4^+^ Foxp3^+^ Treg, there is mounting evidence supporting the existence of a CD8^+^ T regulatory subset, but the precise signals controlling differentiation into this fate are not well understood (46). Using a classical in vitro suppression assay with cells from OT-I Bim^-/-^ > RIP-mOVA chimeras as a source of potential suppressors, we did not observe decreased activation of wildtype OT-I T cells. Additionally, in our co-adoptive transfer experiments, neither the total splenocyte nor the CD8^+^ T cell compartments appeared to contain dominant suppressors. These findings indicate that the functional impairment observed when clonal deletion is prevented is cell-intrinsic in nature. Furthermore, it would suggest that simple impairment of clonal deletion does not lead to the generation of CD8^+^ T regulatory cells and that other factors are involved. However, it remains possible that the mechanisms of tolerance vary with the antigen, quantity of antigen, and presenting APC, as has been previously suggested (8, 9).

It is known that the co-inhibitory receptor PD-1 plays a role in regulating T cell function and is induced in DP thymocytes following strong TCR signaling. In the 2C TCR transgenic model of negative selection, PD-1 deficiency resulted in enhanced generation of DN thymocytes (24) while no such effect was observed in MHCI- or MHCII-restricted male antigen reactive TCR transgenic models (25). We found that PD-1 was similarly induced on OT-I CD8SP thymocytes that survived deletion in RIP-mOVA recipients. Deletion of PD-1 did not dramatically affect the number of OT-I CD8SP thymocytes in RIP-mOVA chimeras, although there were increased numbers of OT-I T cells in the spleen and pancreatic lymph nodes of these mice, as was similarly observed for MOG-specific CD4^+^ T cells on a polyclonal background (47). We think it is unlikely that tolerance was broken due to the elevated numbers of Ag-specific cells, but rather that PD-1 signaling induced a tolerogenic program in the OT-I T cells, since OT-I Bim^-/-^ > RIP-mOVA chimeras contain substantially more OT-I T cells than OT-I PD-1^-/-^ > RIP-mOVA counterparts and tolerance was still enforced in OT-I Bim^-/-^ T cells (6, 7).

PD-1 has primarily been investigated for its role in peripheral tolerance by limiting reactivation, function, and trafficking of self-reactive T cells (10), but a role for PD-1 in establishing tolerance in the thymus has not be reported. Through an adoptive transfer approach using thymocytes from OT-I Bim^-/-^ > RIP-mOVA chimeras as a source of auto-reactive T cells and sub-lethally irradiated PD-L1^-/-^ RIP-mOVA recipients, we found that acute PD-1 signaling during development was sufficient to establish stable tolerance. Furthermore, PD-1 blockade in tolerant OT-I Bim^-/-^ > RIP-mOVA chimeras was unable to break tolerance. This may be due to the blockade treatment falling short of 100 % effectiveness; in this case, even low-level PD-1 signals may filter through and impact tolerance. Additionally, since PD-L1^-/-^ recipients of OT-I Bim^-/-^ > RIP-mOVA thymocytes only began to go diabetic ~ 15 weeks post-adoptive transfer, it may be that PD-1 antibody blockade was too brief to observe a similar effect on tolerance (Fig. 5H). That PD-1 signaling during thymic selection is sufficient to induce tolerance is further supported by Thangavelu *et al*. where PD-1-deficiency in polyclonal thymocytes or recent thymic emigrants, but not established peripheral T cells precipitated a lethal, multi-organ autoimmunity when adoptively transferred into Rag^-/-^ recipients (25). Together with our findings, this suggests our conclusions are broadly applicable to thymic selection overall. It was also recently shown that PD-1 could delay lethal autoimmunity in *Aire*^-/-^ mice (48); in the absence of AIRE in this model, many TRA would not be expressed in the thymus and instead would be first encountered in the periphery, where PD-1 signals could induce protection against autoimmunity. Finally, intestinal inflammation and mortality driven by naïve OT-I T cells in iFABP-OVA mice was shown to be restricted by PD-L1 blockade only when the treatment was administered the day of adoptive transfer, but not 30 days later (49). Given observations made by us and others, we thus hypothesize that it is upon the first high affinity antigen exposure that PD-1 signals are required to induce a durable tolerant state that resists reinvigoration. This is in contrast to previous studies (50–53) proposing only a minor role for PD-1 in early tolerance induction (see review (54)). However, many of these studies were performed on the NOD background, wherein native tolerance is not established (55).

While acute PD-1 signals received by OT-I T cells in the thymus provided a relatively durable tolerance when OT-I thymocytes were adoptively transferred into sub-lethally irradiated RIP-mOVA PD-L1^-/-^ recipients, we cannot formally discount a role for PD-L2. However, it appears that PD-L2 does not exert a large effect on tolerance in this model. In our studies with OT-I Bim^-/-^ > RIP-mOVA PD-L1^-/-^ chimeras, tolerance is not established, and diabetes is induced rapidly even with unaltered PD-L2 expression. Similarly, the Sharpe group previously demonstrated in the NOD background a unique role for PD-L1 in restraining self-reactive T cells in the pancreas (56). Additionally, thymocytes or splenocytes from OT-I Bim^-/-^ > RIP-mOVA mice adoptively transferred into RIP-mOVA PD-L1^-/-^ recipients did not maintain tolerance indefinitely. Since in PD-L1 sufficient recipients adoptively transferred thymocytes remained tolerant throughout the entire course of the experiment, PD-1 signaling induced via interactions with PD-L1 likely aided in the maintenance of tolerance. It will be interesting to assess the necessity for thymic PD-1 signaling by limiting PD-1 interactions to the periphery (e.g adoptive transfer of thymocytes from OT-I Bim^-/-^ > RIP-mOVA PD-L1^-/-^ chimeras into PD-L1^+/+^ recipients), in which case failure to establish tolerance would further suggest it is PD-1 signals received specifically during first encounter with Ag in the thymus that are necessary for establishing tolerance. Indeed, recent thymic emigrants are thought uniquely sensitive to tolerizing factors compared to more mature circulating T cells, suggesting that PD-1 signals delivered in the thymus and periphery may lead to functionally different outcomes (57).

Work done by the Fife group showed that established anergic peripheral T cells are not reinvigorated by PD-L1 blockade (49). They and others postulated that this is due to an epigenetic landscape resistant to reinvigoration in the context of “exhaustion” and tolerance (58–61). Given this, we hypothesize that acute PD-1 signaling supports long-term tolerance by inducing a tolerant epigenetic memory. Using a double transgenic TCR_GAG_ Alb:GAG model, the Greenberg group recently showed that self-reactive T cells that escape thymic deletion do not cause liver injury or respond to immunization with LM-GAG. These cells regained function through robust lymphopenia-induced proliferation (LIP) and responded to LM-GAG immunization at early time points, but 3-4 months post-adoptive transfer, functional impairment was reacquired. Their observations were similarly recorded following transfer into both high affinity Ag-bearing (Alb:GAG) and Ag-lacking (C57BL/6) recipients (58). Conversely, we failed to observe LIP by the functionally impaired CD8^+^ T cells from OT-I Bim^-/-^ > RIP-mOVA chimeras and instead observed a dependence on persistent high affinity antigen for this functional impairment. This difference may be because OT-I thymocytes developing in a RIP-mOVA thymus normally undergo efficient thymic deletion, while in the TCR_GAG_ Alb:GAG model, deletion is only partial. Further, it is unclear whether the functional impairment observed in TCR_GAG_ T cells was established in the thymus, as was the case for OT-I T cells; therefore, the states of dysfunction our groups documented may not be analogous (58). The tolerance-specific epigenetic program they and others propose drives tolerance in self-reactive T cells (58, 59) may also underlie our observations. This phenomenon is a clear obstacle for immunotherapies seeking to reinvigorate functionally impaired T cells, thus future investigation should aim to determine the mechanism by which this occurs and its relationship with PD-1.

Immune checkpoint blockade (ICB) therapies targeting the PD-1:PD-L1 signaling axis have received considerable attention in the last 10 years for their effectiveness in treating a number of different cancers. A recent study proposed that ICB targeting the PD-1:PD-L1 signaling axis promoted the activation and expansion of newly recruited clones to an anti-tumour response, rather than reinvigorating pre-existing tumour-infiltrating T lymphocytes as is often suggested to underlie the success of PD-1 blockade (62). In this study, we provide evidence to support this alternate mechanism of anti-PD-1 ICB therapies. If PD-1 is indeed critical for the establishment of tolerance rather than its maintenance as we suggest, it follows that PD-1-targeting ICB may enhance recruitment of clones naïve to the PD-1 tolerogenic signal. Targets of these therapies may originate in the periphery as tumour-specific naïve T cells whose primary encounter with antigen is no longer rendered tolerogenic by PD-1, or in the thymus as tumour-harboured self-antigen-specific thymocytes whose native tolerance through PD-1 is prevented. Reinvigoration of “exhausted” CD8 T cells is still possible through processes independent from what we have observed in this study, but here we introduce new considerations for the understanding of anti-PD-1 targeting blockade therapies.

Despite the success of ICB therapies, clinical responses vary with different types of cancer, and immune-related adverse events (IRAE) are a persistent risk (11, 63). It is conceivable that should PD-1 blocking antibodies reach the thymus, normal selection processes may be hindered. While this provides a potential explanation for the development of some IRAEs, it may also be in some cases the mechanism behind successful cancer immunotherapies. This highlights the need for a greater understanding of the mechanism by which PD-1 supports normal tolerance and how this relates to ICB therapies. Overall, we provide evidence for a novel role for PD-1 in the establishment of tolerance to TRA in the thymus in escapees from clonal deletion. Depending on the conditions of antigen presentation (antigen density, presenting cells, location of first encounter), PD-1 signals may induce states of tolerance or simply aid in its maintenance, potentially with different functional outcomes.

## Acknowledgements

The authors thank Dr. Heather Melichar and Stefanie Valbon for critical review of the manuscript. Flow cytometry experiments were performed at The University of Alberta Faculty of Medicine & Dentistry Flow Cytometry Facility, which receives financial support from the Faculty of Medicine & Dentistry and Canada Foundation for Innovation (CFI) awards to contributing investigators. Health Sciences Laboratory Animal Services and Mr. Bing Zhang provided animal husbandry and technical assistance.

## Footnotes

- This work was supported by a grant from the Canadian Institutes of Health Research (PS 156104) to T.A.B.

## Abbreviations used in this article

WT: *Wildtype*
TRA: Tissue-Restricted Antigen
UbA: Ubiquitous Antigen
pMHC: peptide-MHC
BM: Bone Marrow

