## Supplemental Materials for "Establishment of CD8^+^ T cell thymic central tolerance to tissue-restricted antigen requires PD-1"

**Supplemental figures**

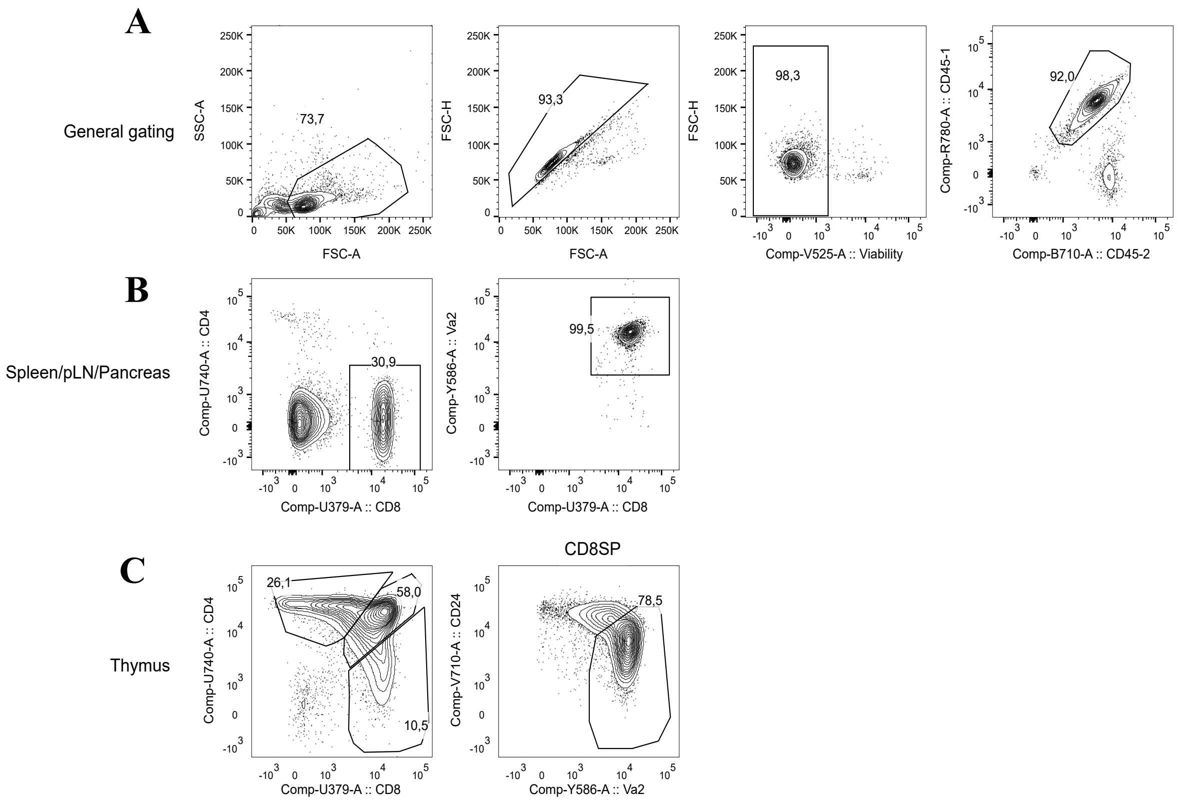


**S1**. Gating strategy for analysis of OT-I transgenic thymic and peripheral T cells. Gates are ordered hierarchically left to right. A, General gating strategy at the beginning of all analyses. B, Gating strategy for identifying peripheral OT-I T cells. C, Gating strategy for identifying OT-I thymocytes; the mature thymocyte gate is drawn within the CD8SP population.

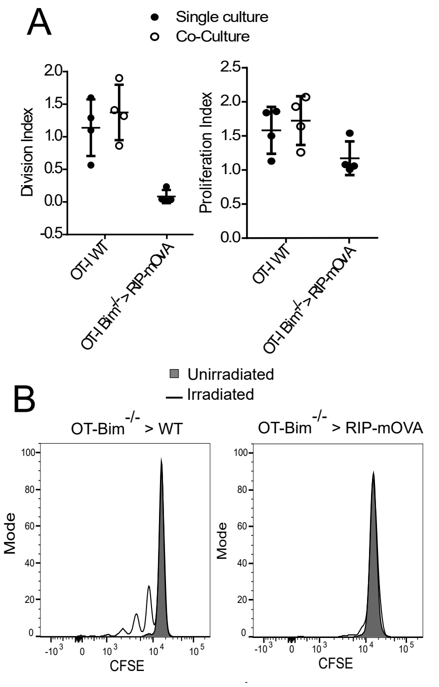


**S2**. A, Day 2 proliferation from experiments represented in Figure 1. B, Lymphopenia induced proliferation of $V\alpha 2^{+}$ $CD8^{+}$ splenocytes transferred from OT-$Bim^{-/-}$ > WT or OT-I $Bim^{-/-}$ > RIP-mOVA chimeras into unirradiated (grey) or sub-lethally irradiated (open) recipients, measured by CFSE labelling (n = 2). For each assay described above, the number of independent experiments (IE) ≥ 2. Data are mean ± SD.

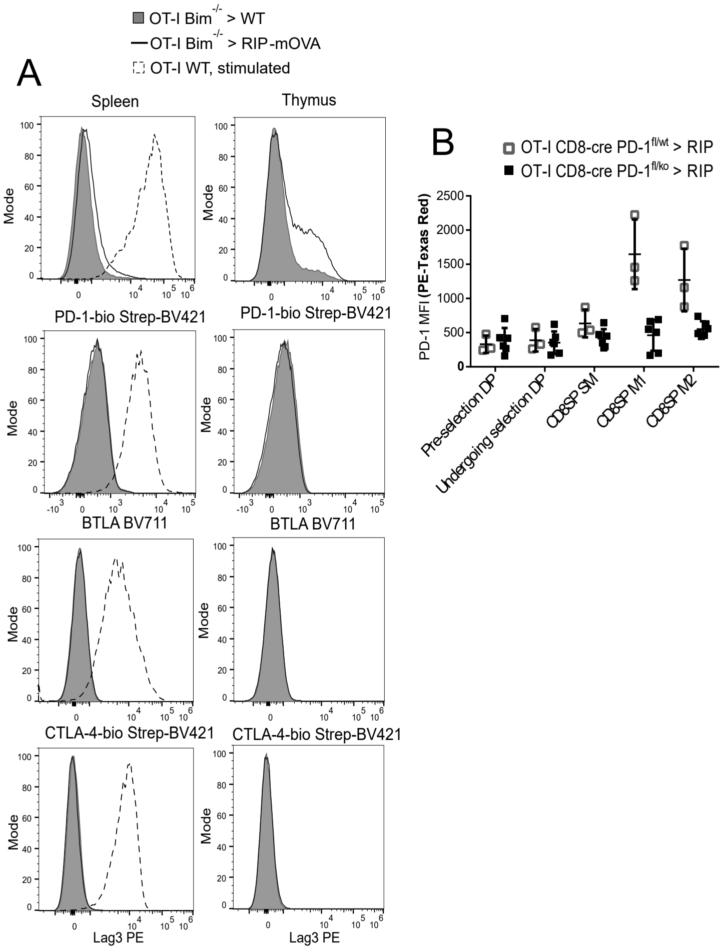


**S3.** A, Co-inhibitory molecule expression in OT-I Bim$^{-/-}$ > RIP-mOVA and OT-I Bim$^{-/-}$ > WT chimeras. As a positive control for co-inhibitory molecule expression, WT OT-I splenocytes were stimulated for 24 hr with high affinity peptide OVA (100 nM). B, PD-1 expression in RIP-mOVA chimeras with OT-I CD8-cre PD-1$^{fl/wt}$ or PD-1$^{fl/ko}$ bone marrow. PD-1 expression was analyzed within the PD-1$^{+}$ subset within each population shown. Pre-selection DP thymocytes (CD69$^{lo}$ TCRβ$^{lo}$), DP undergoing selection (CD69$^{hi}$ TCRβ$^{hi}$), SM thymocytes (TCRβ$^{+}$ H-2K$^{b\ hi}$), M1 thymocytes (TCRβ$^{+}$ H-2K$^{b\ hi}$ CD69$^{hi}$), M2 thymocytes (TCRβ$^{+}$ H-2K$^{b\ hi}$ CD69$^{hi}$) (n = 4). For each assay described above, the number of independent experiments (IE) ≥ 4.

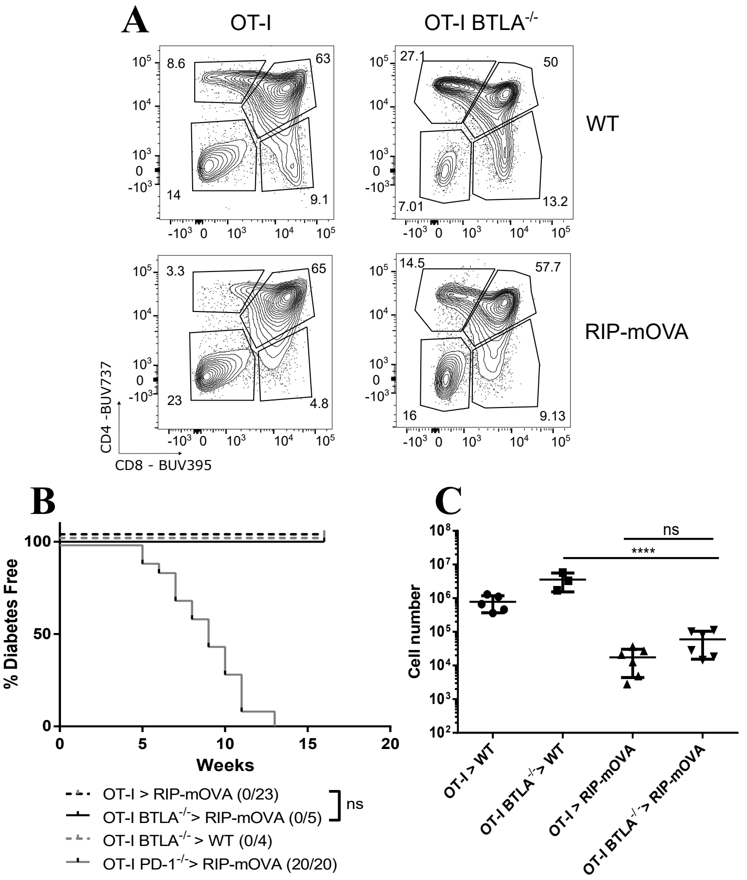


**S4**. A, Thymic profiles of WT or BTLA$^{-/-}$ OT-I > RIP or B6 chimeras. B, Incidence of diabetes in RIP-mOVA or WT chimeras with BTLA$^{-/-}$ (IE = 2) or WT bone marrow. C, Number of mature V$\alpha$2$^{+}$ CD8SP CD24$^{lo}$ thymocytes in the chimeras described in (A); OT-I > WT (n = 5), OT-I BTLA$^{-/-}$ > WT (n = 3), OT-I > RIP-mOVA (n = 6), OT-I BTLA$^{-/-}$ > RIP-mOVA (n = 6). Data are mean ± SD. Asterisks represent statistical significance as determined by Log-rank test (B) or two-way ANOVA with Sidak's multiple comparisons tests (C). * $p \le 0.05$, ** $p \le 0.01$, *** $p \le 0.001$, **** $p \le 0.0001$.
